## Supplementary Figure 1 for "GLP-1R associates with VAPB and SPHKAP at ERMCSs to regulate β-cell mitochondrial remodelling and function"

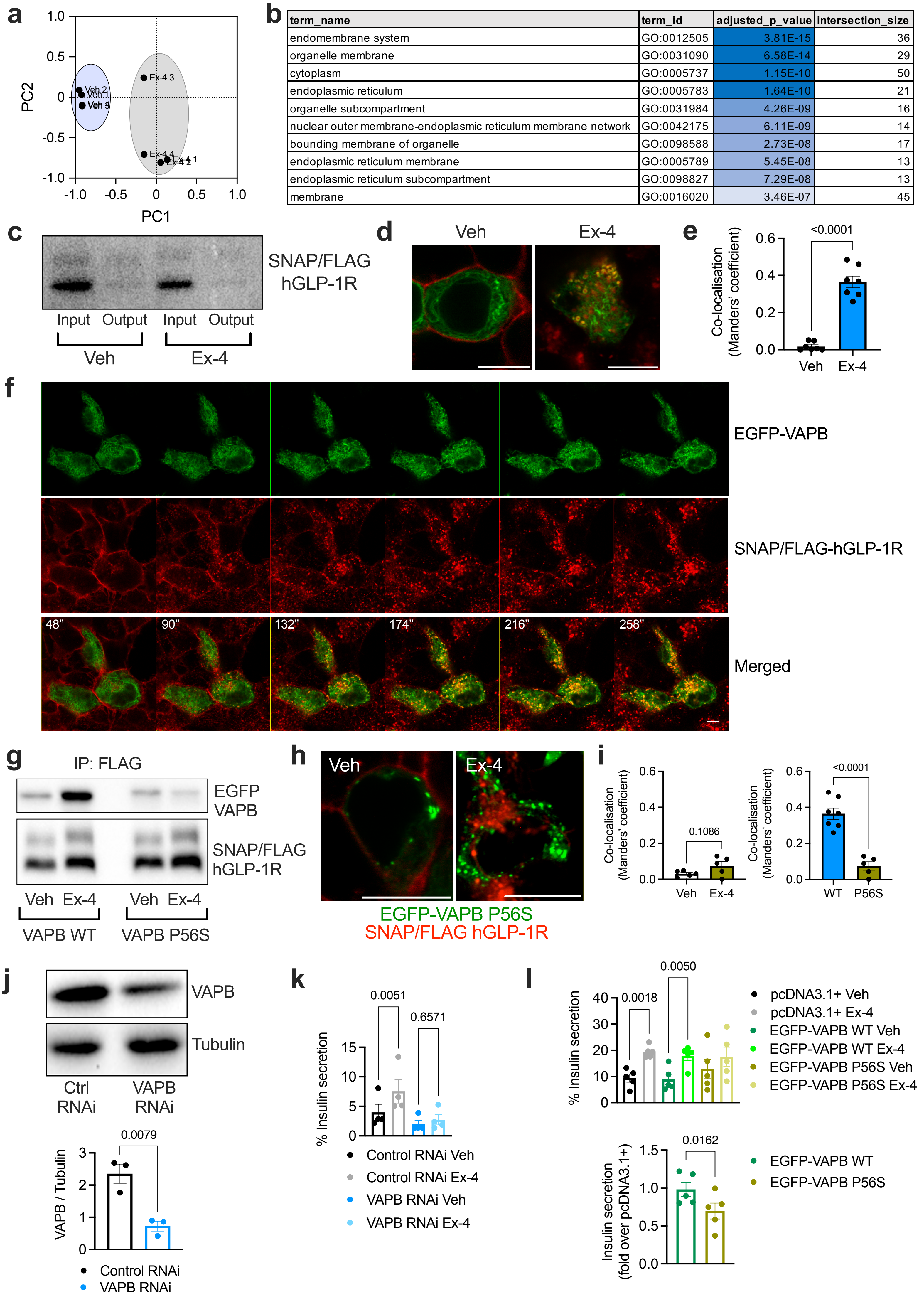


| term_name | term_id | adjusted_p_value | intersection_size |
|---|---|---|---|
| endomembrane system | GO:0012505 | 3.81E-15 | 36 |
| organelle membrane | GO:0031090 | 6.58E-14 | 29 |
| cytoplasm | GO:0005737 | 1.15E-10 | 50 |
| endoplasmic reticulum | GO:0005783 | 1.64E-10 | 21 |
| organelle subcompartment | GO:0031984 | 4.26E-09 | 16 |
| nuclear outer membrane-endoplasmic reticulum membrane network | GO:0042175 | 6.11E-09 | 14 |
| bounding membrane of organelle | GO:0098588 | 2.73E-08 | 17 |
| endoplasmic reticulum membrane | GO:0005789 | 5.45E-08 | 13 |
| endoplasmic reticulum subcompartment | GO:0098827 | 7.29E-08 | 13 |
| membrane | GO:0016020 | 3.46E-07 | 45 |
