## Supplementary figures and images for "GLP-1R associates with VAPB and SPHKAP at ERMCSs to regulate β-cell mitochondrial remodelling and function"

### Supplementary Figure 2

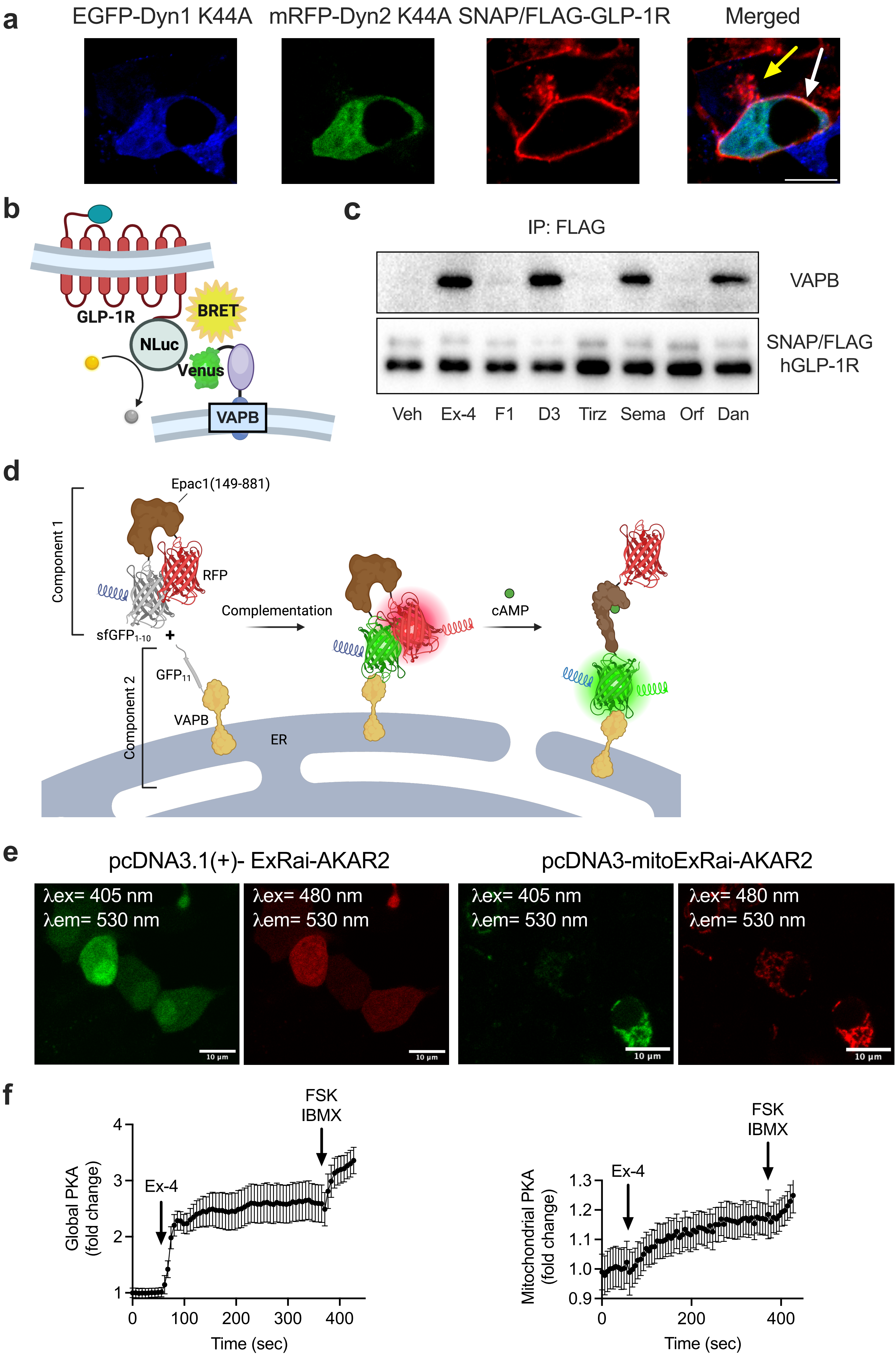

### Supplementary Figure 3

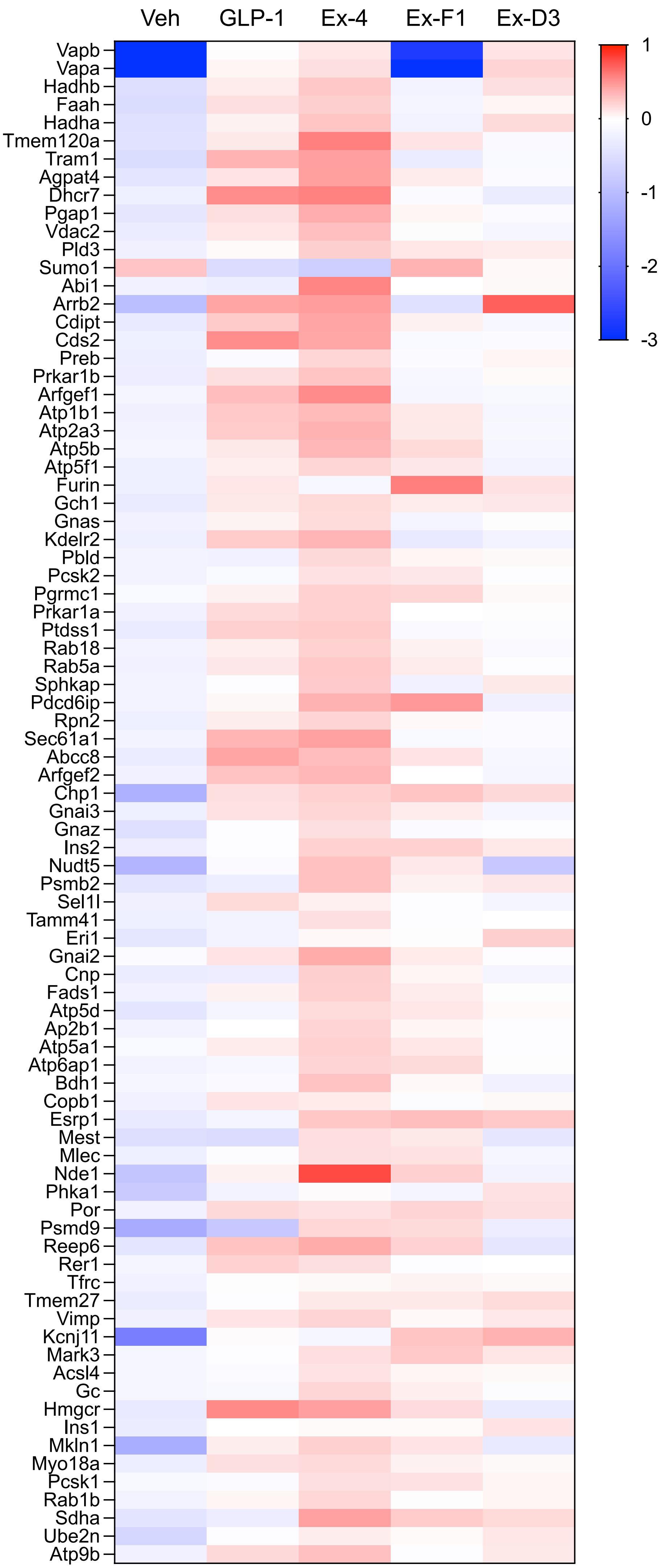

### Supplementary Figure 4

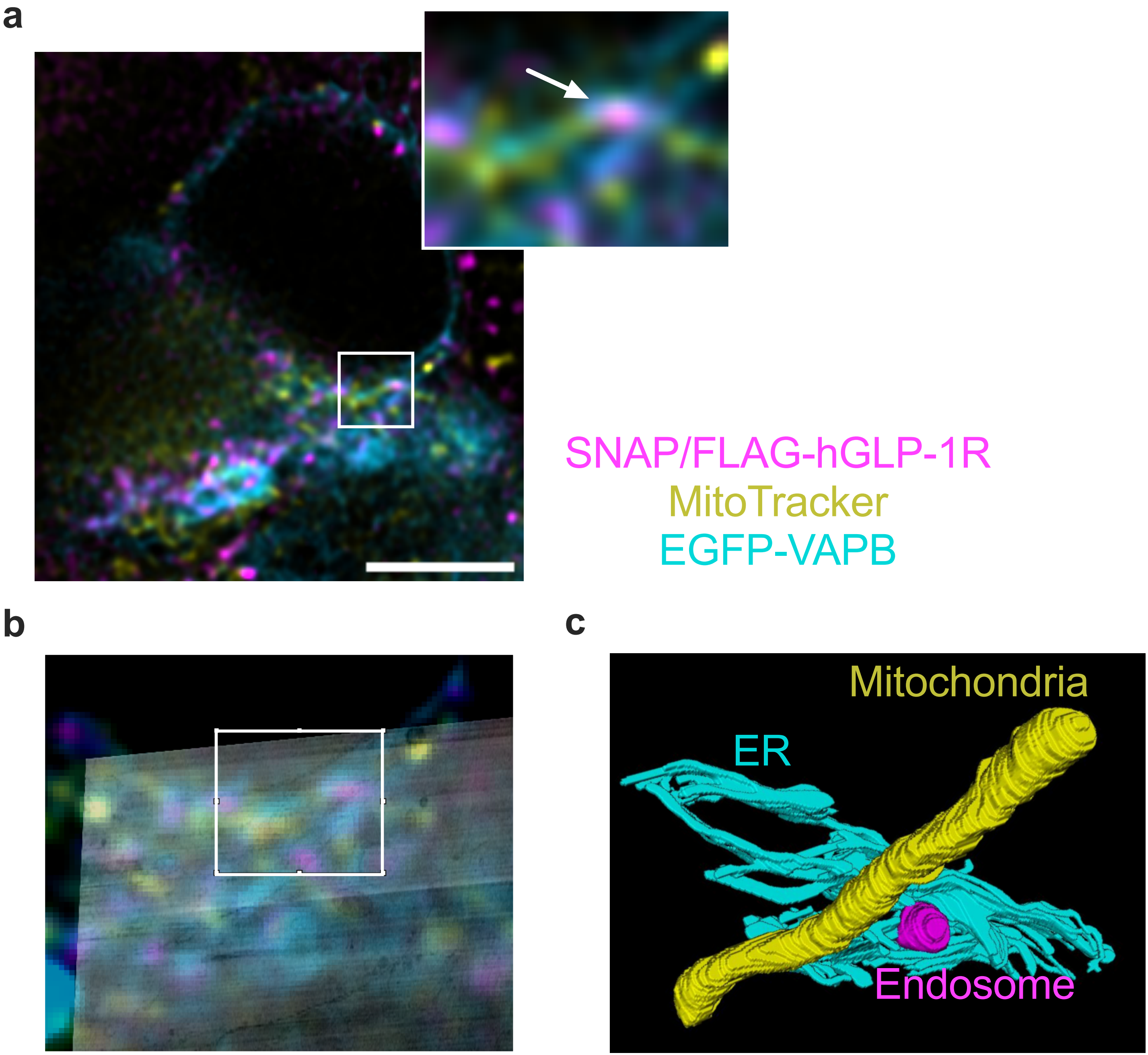

### Supplementary Figure 5

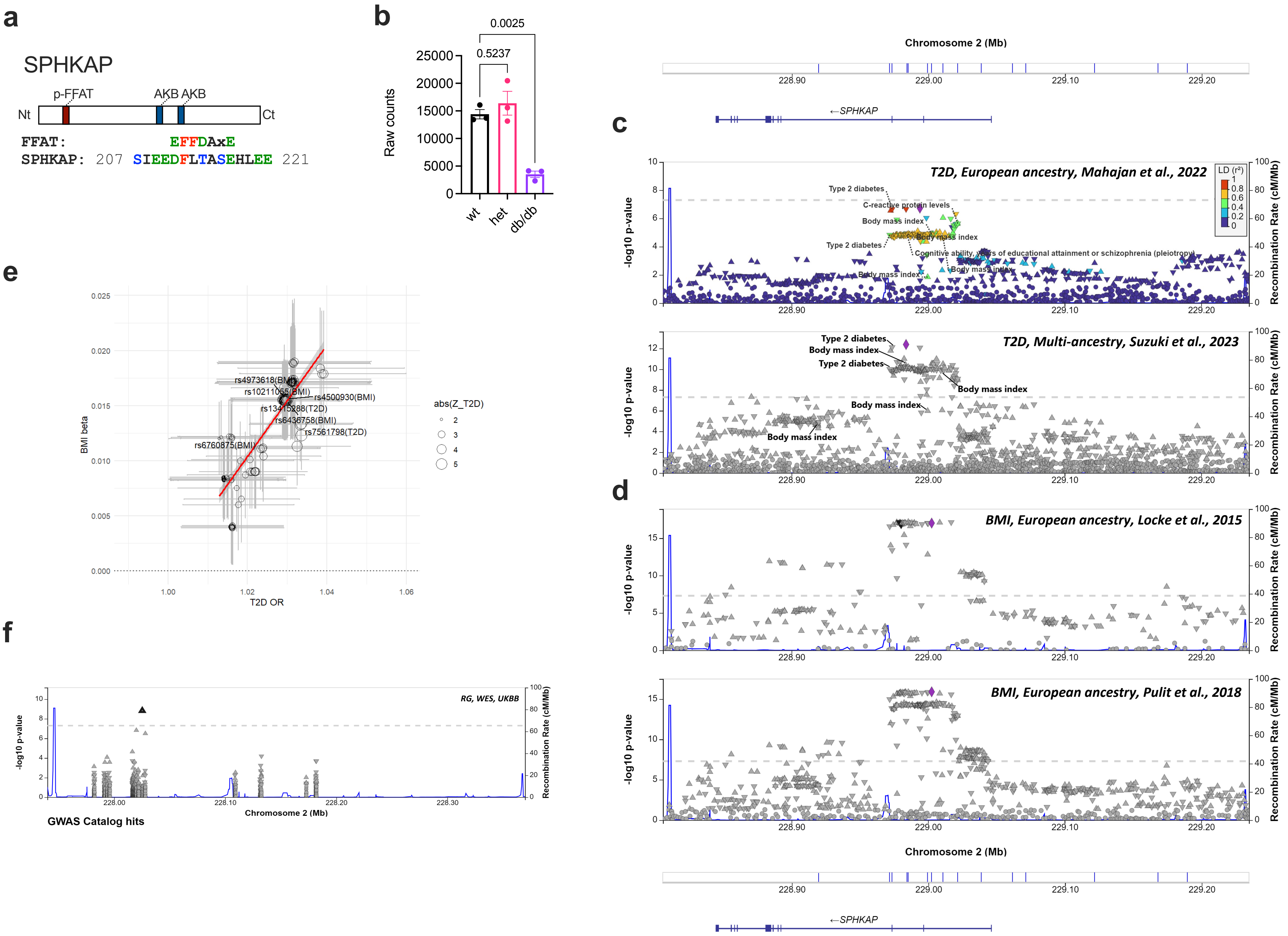

### Supplementary Figure 6

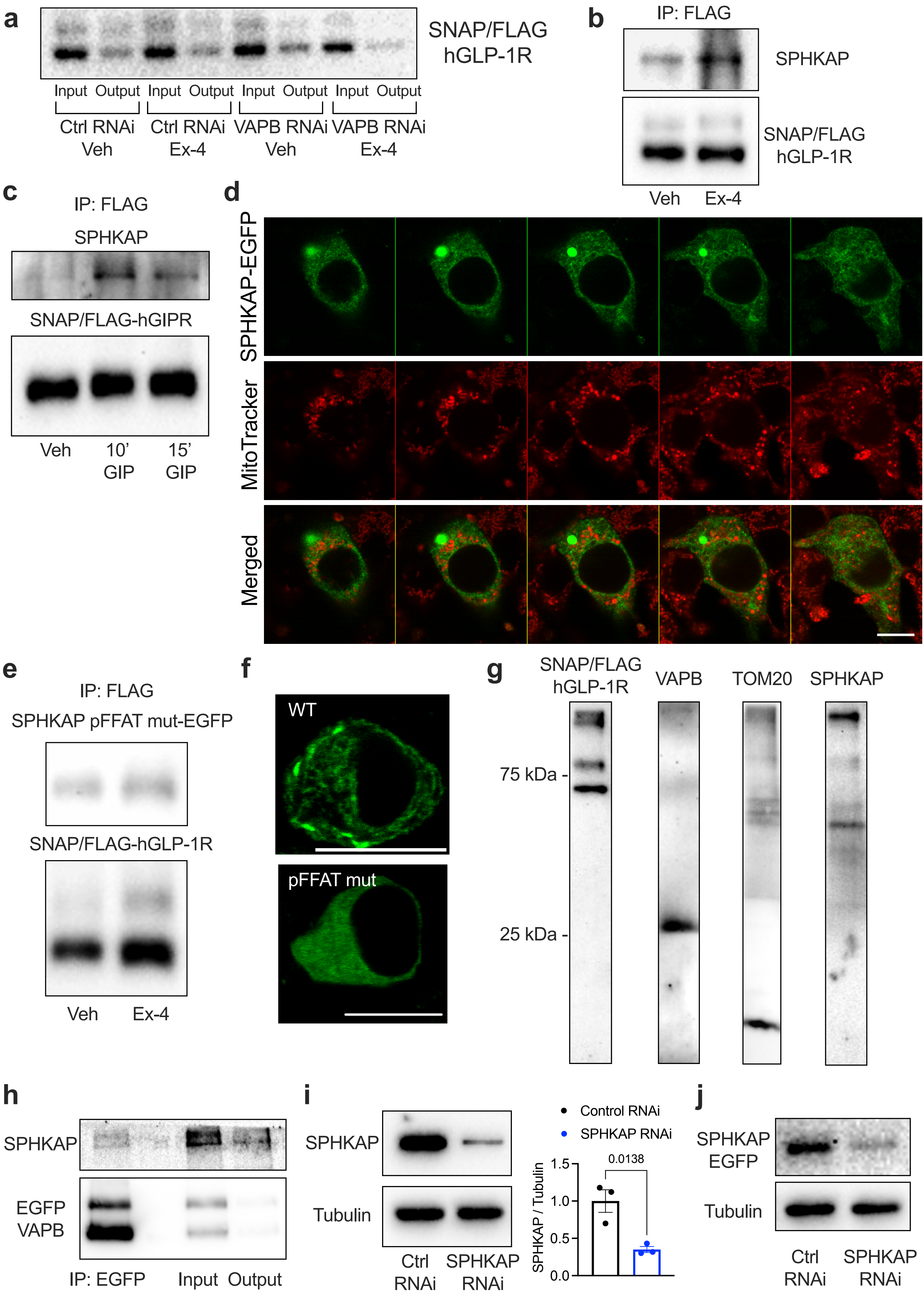

### Supplementary Figure 7

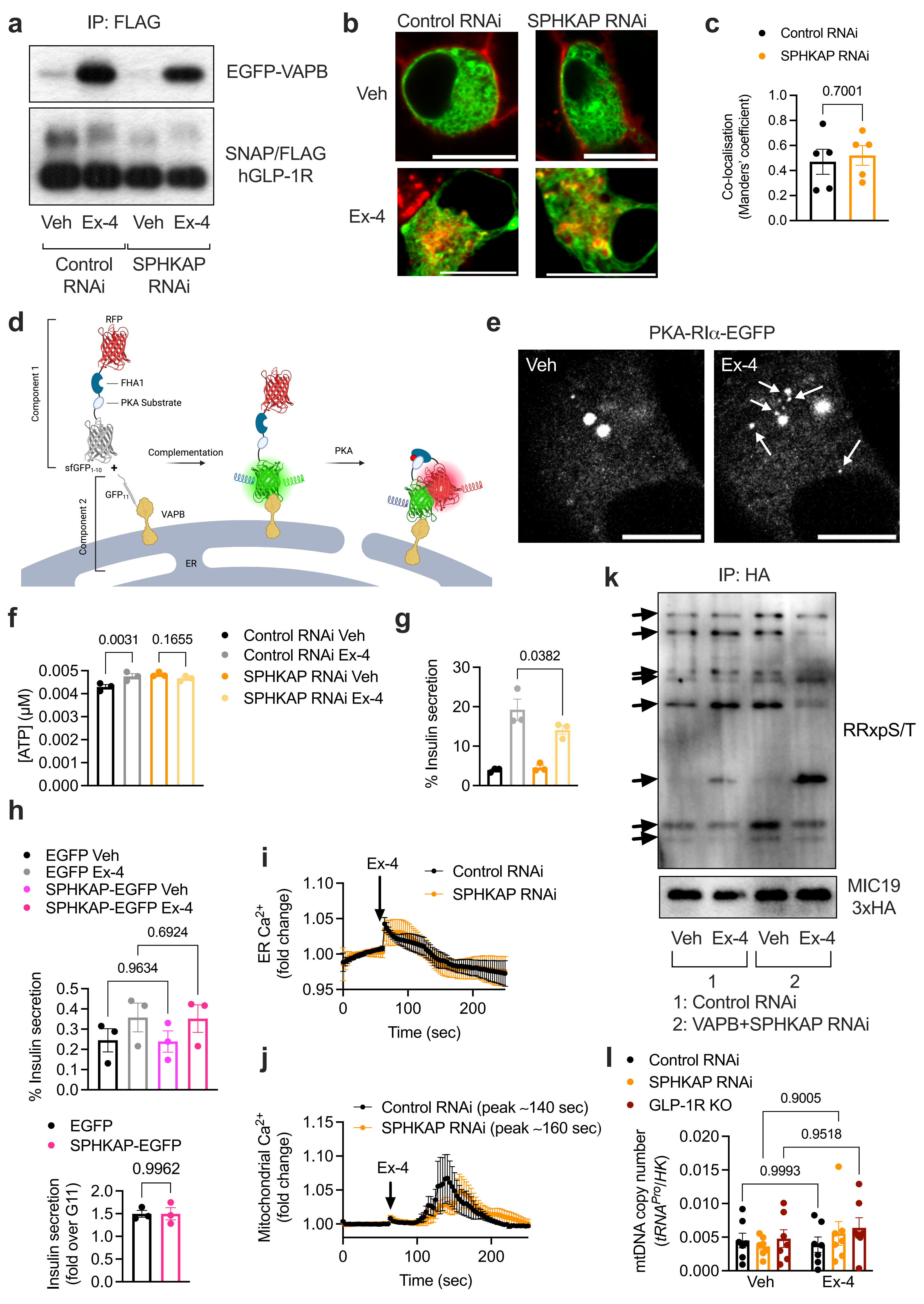

### Supplementary Figure 8

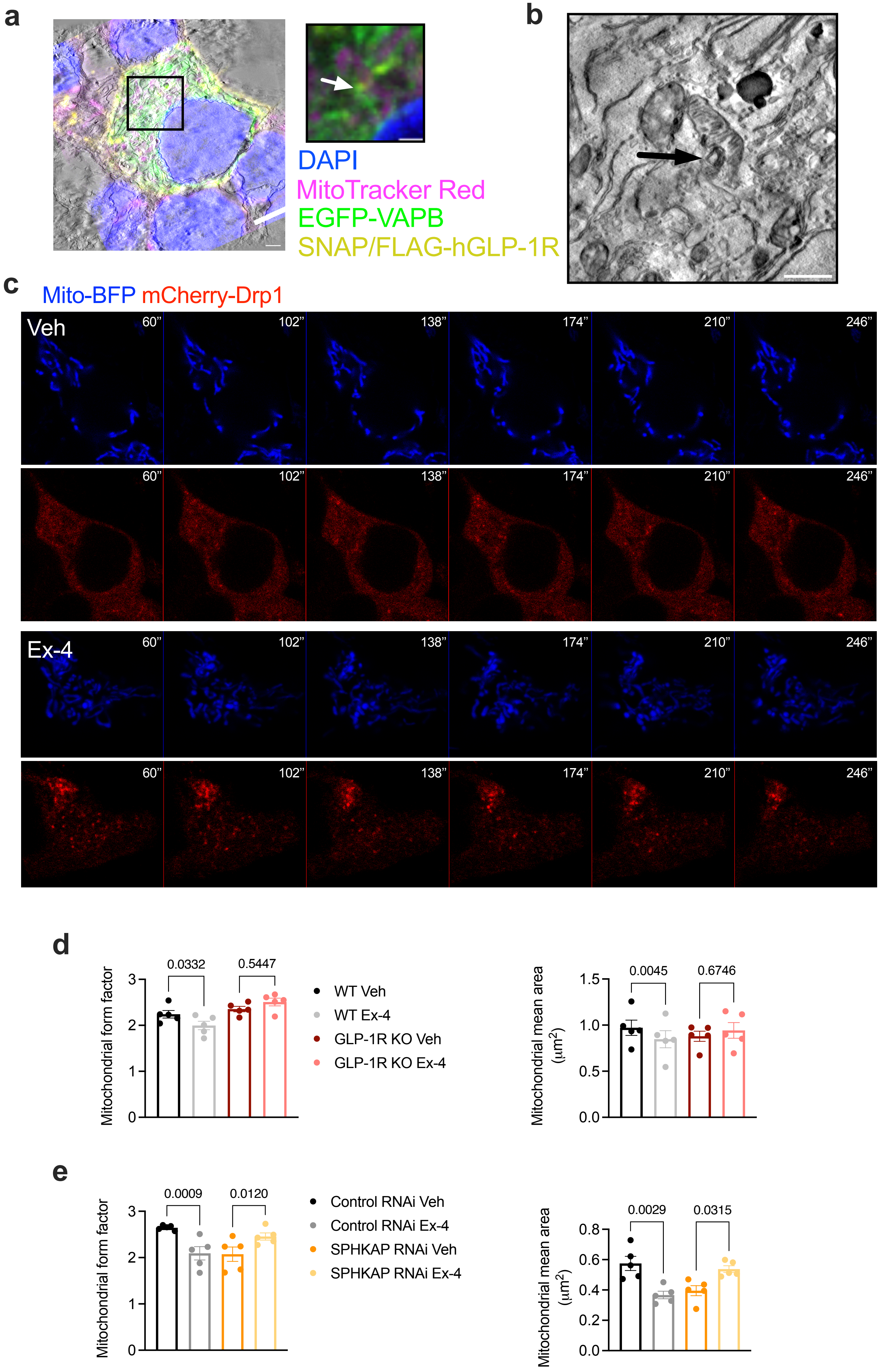

### Supplementary Figure 9

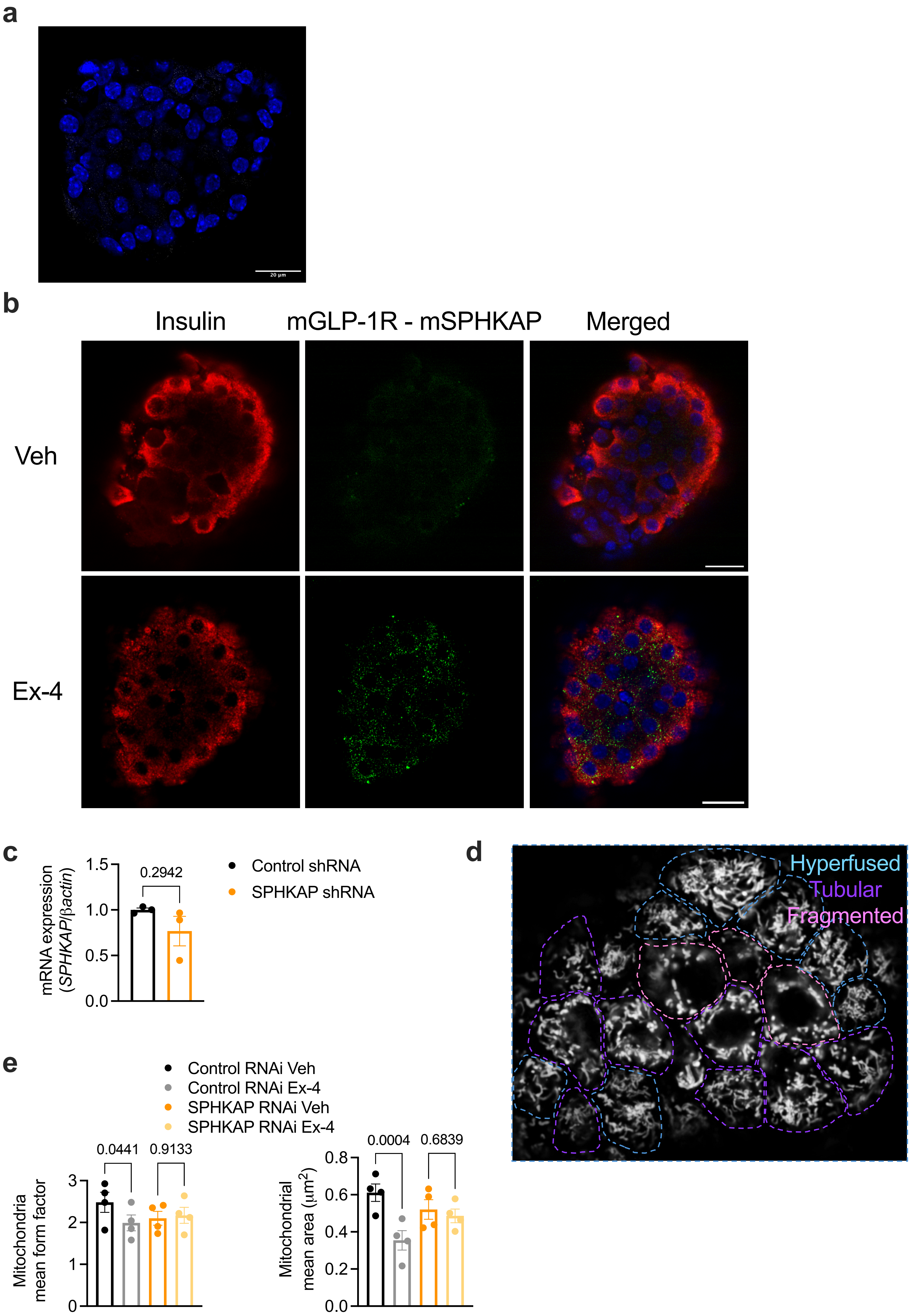
